# Highly frequent undesired insertional mutagenesis during *Drosophila* genome editing

**DOI:** 10.1101/2025.08.11.669658

**Authors:** Emma Källstig, Evelyne Ruchti, Medha Raman, Jamshid Asadzadeh, Bernard L. Schneider, Brian D. McCabe

## Abstract

CRISPR/Cas9 based genome editing employing Homology Directed Repair (HDR) from template vector sequences is a widely used technique to enable precise insertions, deletions or modifications to genes. Here, we describe an undesired and highly frequent editing event when using conventional CRISPR/Cas9 plus HDR methods for *Drosophila melanogaster* germline genome editing. We find that the template vector employed for HDR repair unwantedly and commonly inserts into the genome. We observe this deviation from the desired edit at multiple genomic locations, with different HDR vectors and with multiple genome editing designs. To avoid these events, we have generated a novel HDR template vector that enables animals with these undesired insertions to be identified and excluded. Our results suggest that HDR based genome edited animals must be carefully screened for unwanted vector template genomic integration in order to avoid misleading interpretations of genome editing outcomes.

## Introduction

Genome engineering in model organisms and cell lines is a rapidly evolving field with continuously developing methods, techniques, and approaches (1, 2). The field was revolutionized by the employment of the clustered regularly interspaced short palindromic repeat (CRISPR)/Cas9 system, derived from prokaryotic immunity, which enables the precise targeting and subsequent modification of genes (3). The system forms the basis to create many types of genetic modifications, for example, insertions, deletions, or substitutions in genes of interest, with improved ease, speed, and accuracy (4). CRISPR/Cas9 based genome editing relies upon guide RNAs (gRNAs) to direct the endonuclease Cas9 protein to a target DNA sequence, leading to the introduction of double-stranded DNA breaks (DSBs) (5). The cell can then repair these DSBs by either nonhomologous end joining (NHEJ) or homology-directed repair (HDR). NHEJ ligates nonhomologous DNA ends, which if occurring erroneously, can induce either insertions or deletions (indels) (6). HDR on the other hand, uses a DNA template from a homologous DNA strand to repair DSBs (7). Providing an exogenous ‘template’ sequence for HDR, commonly delivered from a plasmid vector, enables exploitation of the HDR process to create specific genome insertions, deletions or other nucleotide modifications, when these changes are introduced into the exogenous HDR template. The adaptability of the approach has enhanced the popularity of CRISPR/Cas9 as a technique for germline genome editing in model organisms (8) including *Drosophila melanogaster* (9). Germline genome editing in *Drosophila* employing HDR is most frequently carried out by germline injection of a HDR template plasmid, which contains homology arms flanking the desired edited sequence, together with plasmids expressing one or more gRNAs designed to induce DSBs in the targeted locus. To induce recombination, Cas9 is introduced in the germline either via transient expression from a plasmid or using transgenic animals stably expressing Cas9 (Figure 1) (10). Successfully edited animals can then be identified by molecular screening (e.g. PCR), phenotypic alterations or selectable markers (10).

**Figure 1.**
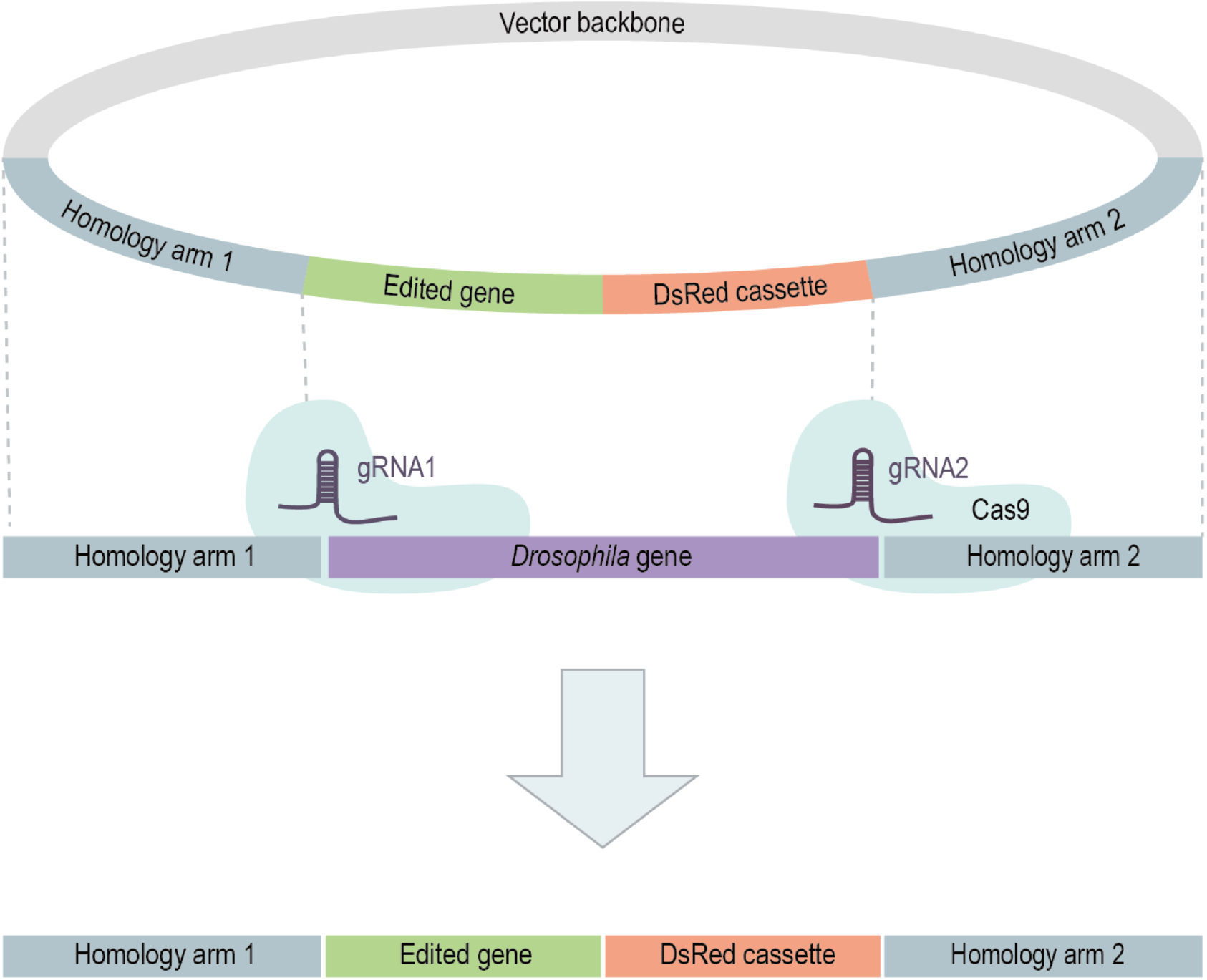
Schematic of the use of CRISPR/Cas9 HDR strategy. *Drosophila* embryos, expressing germline Cas9, are injected with plasmids expressing two gRNAs in addition to a plasmid containing the edited target sequence flanked by *∼*1 kb long homology arms which act as a template for HDR. An eye expression DsRed selectable cassette is also included to facilitate identification of edited animals. After the cleavage of the targeted locus by gRNA/Cas9, homology to the exogenous template sequence enables HDR, resulting in the conversion of the genomic sequence to match the template plasmid sequence.

While there are many advantages to the use of CRISPR/Cas9 based genome editing compared to other available methods, several studies have also revealed drawbacks of the technique. In mouse embryonic stem cells, hematopoietic progenitors and human differentiated cell lines, the DSBs created by guide RNAs and Cas9 have the potential to induce large deletions or rearrangements of the genomes (11). Similar undesired large deletions and insertions have been observed during mouse zygote genome editing (12). Here, we describe an undesired and highly frequent mutagenic insertion of the HDR donor plasmid template vector into the genome. Analogous gene editing problems have been previously identified (13–17), however, the frequency of these undesired events was not fully apparent. Here, we observe that HDR vector template insertion occurs to an alarmingly high degree with multiple template vectors. Our results caution investigators carrying out HDR directed genome editing in *Drosophila* to be on alert for erroneous template genomic insertion and mutagenesis, which may obscure and confound analysis of genome edited animals. To combat this problem, we have designed and tested a novel and versatile HDR template vector that allows the identification and exclusion of these undesired mutagenic insertional events, in addition to enabling ‘all-in-one’ template plus gRNA sequences single vector HDR genome editing.

## Results

### Undesired insertional mutagenesis induced during the generation of genome edited humanised *Drosophila*

The TAR DNA-binding protein 43 (*hTDP-43*) is associated with both sporadic and familial Amyotrophic lateral sclerosis (ALS) (18). With the goal to generate high fidelity *Drosophila* models of *hTDP-43* ALS, we sought to create Human Gene Replacement (HGR) or ‘humanised’ genome edited *Drosophila* lines, where we replaced the *Drosophila* TAR DNA-binding protein-43 homolog (*TBPH*) (19) gene with a human *hTDP-43* encoding cDNA (*hTDP-43*) (20). We began by using CRISPR Optimal Target Finder software (21), to design two gRNAs outside of the *TBPH* 3’ and 5’ UTR regions to induce Cas9 dependent DSBs. Both of the gRNA sequences we designed were synthesised and inserted into pCFD3 to facilitate expression in germline cells by transgene injection. To enable the replacement of the *Drosophila TBPH* open reading frame (ORF) with a human *hTDP-43* ORF while maintaining *Drosophila* 5’ and 3’ regulatory regions, we generated a HDR template plasmid containing ∼1 kb of 5’ and 3’ sequences homologous to the genomic region surrounding *Drosophila TBPH*, together with an artificial replacement exon encoding *hTDP-43*, and a selection cassette to allow identification of successfully edited animals by eye specific DsRed expression (Figure 2A) (22, 23). These sequences were cloned into the plasmid vector pCR4 (24). Both the gRNA encoding plasmids and the HDR replacement cassette were injected into *Drosophila* strains expressing Cas9 in the germline (25). Animals where genome editing had occurred were then identified by eye expression of DsRed. Other than the aim of *Drosophila* to human gene replacement, the design of both plasmids encoding the targeting gRNAs and containing the HDR template followed conventional practices and widely utilised approaches (9, 10).

**Figure 2.**
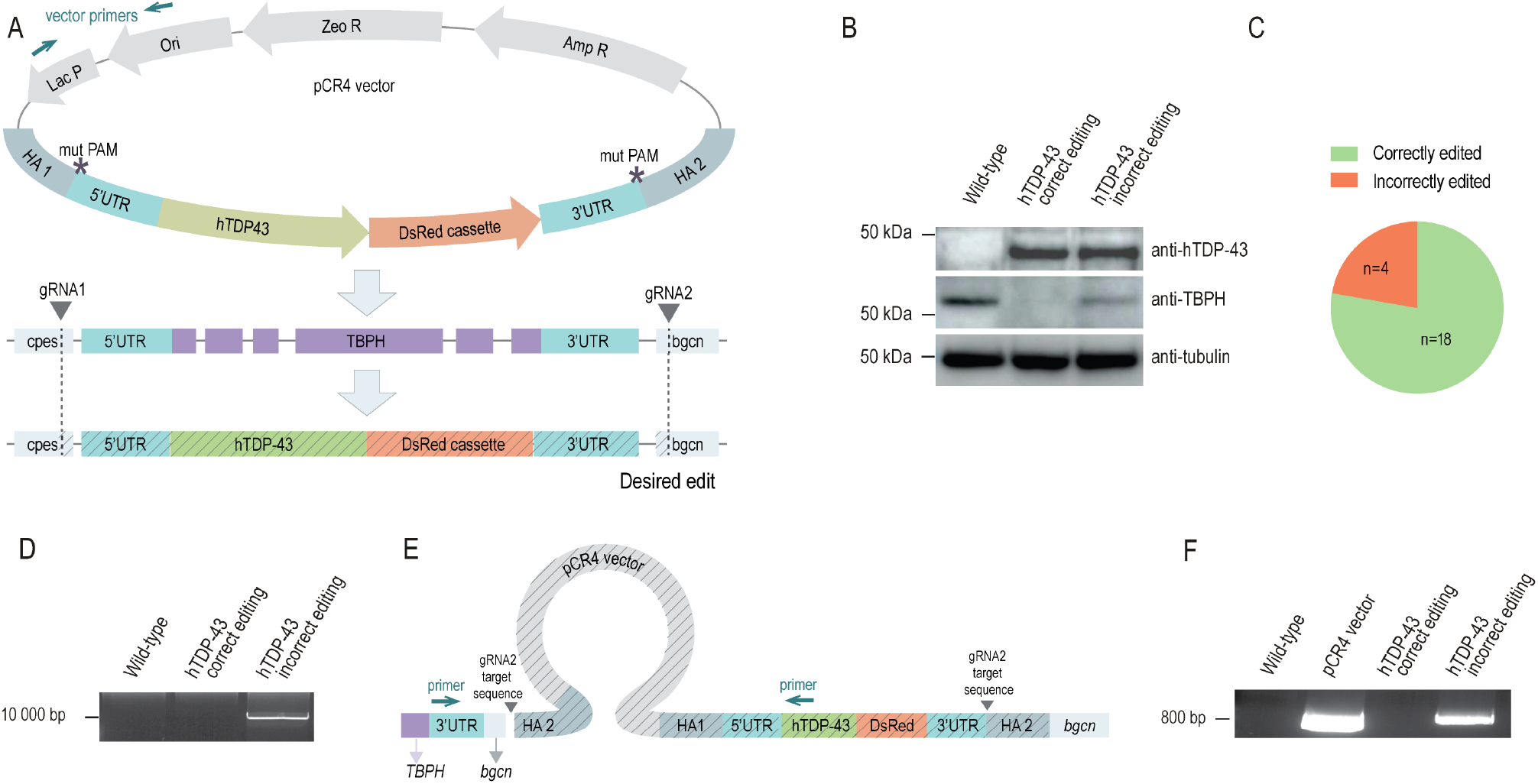
Undesired genomic insertion of HDR template vector sequence during gene replacement HDR. (A) Illustration of the CRISPR/Cas9 HDR replacement of the endogenous *Drosophila TBPH* gene by *hTDP-43*. A HDR template cloned into a circular pCR4 vector includes two *∼*1 kb long homology arms on each side of *hTDP-43*, mutated gRNA PAM sequences [*] and a selectable DsRed cassette. Two gRNAs targeting sequences in the *TBPH* neighbouring genes, *bgcn* and *cpes*, enable Cas9 induced DSBs. The desired HDR editing result is a complete substitution of the *Drosophila TBPH* gene by *hTDP-43* plus insertion of the DsRed cassette. (B) Western blot showing the expression of hTDP-43, *Drosophila* TBPH and tubulin loading control in protein extracts derived from *Drosophila* controls, correctly edited humanised *hTDP-43 Drosophila*, and a humanised *hTDP-43 Drosophila* line with incorrect editing due to template genomic insertion. (C) Chart illustrating the fraction of humanised *hTDP-43 Drosophila* lines that resulted in an incorrect insertion of the template vector backbone (n=4), compared to the fraction of lines with *TBPH* correctly replaced by *hTDP-43* (n=18). (D) Image of an agarose gel from a genomic DNA PCR using a forward primer in the *TBPH* 3’ end and a reverse primer in the *hTDP-43* 5’ end, indicated in panel (E), which produces a product of *∼*10 kb from genomic DNA derived from incorrectly edited *hTDP-43 Drosophila*, which is not present in genomic DNA from *Drosophila* controls or a correctly edited replacement *hTDP-43* line. (E) Schematic of the integrated vector sequence in incorrectly edited *hTDP-43* animals. The entire HDR template vector sequence is inserted into the genome and flanked by sequences matching homology arm 2. Primers used to detect the presence of the undesired vector backbone in the genome are represented by blue arrows. (F) Image of an agarose gel from genomic DNA amplification using primers specific for the template pCR4 vector. Vector primer positions are indicated in panel (A). Vector plasmid sequences can be detected in vector DNA and genomic DNA derived from incorrectly edited *hTDP-43* lines, but not in genomic DNA derived from correctly edited *hTDP-43 Drosophila* or controls. Diagrams are not to scale.

After identification of genome edited animals, individual lines were produced, and homozygous animals from each line were analysed for expression of the *Drosophila TBPH* and *hTDP-43* proteins. Our expectation was that animals would express only *hTDP-43* but not *Drosophila TBPH* if the editing event was successful. Indeed, using western blotting, we did identify 18 of 22 lines where only *hTDP-43* was expressed and *Drosophila TBPH* was undetectable. We subsequently confirmed in these lines using genomic DNA sequencing that the expected editing event had occurred.

However, we found that 18% (4 out of 22) of the lines we generated expressed both *Drosophila TBPH* and human *hTDP-43* (Figure 2B,C). To understand how this unwanted outcome could occur we investigated these lines further. We began by designing a forward primer in the *Drosophila TBPH* gene and a reverse primer for the *hTDP-43* gene in an attempt to amplify the region between these genes. Using genomic DNA generated from *Drosophila* lines expressing both *hTDP-43* and *TBPH* as a template, we amplified a ∼ 10 kb product that was not present in unedited animals or in animals where the editing event had occurred correctly (Figure 2D). We then sequenced this product and discovered that, in addition to including genomic sequences flanking *Drosophila TBPH*, it also included the entire pCR4 plasmid vector sequence, indicating that the entire pCR4 HDR editing template plasmid vector had inserted into the genome of these animals (Figure 2E). This insertion occurred proximal to *Drosophila TBPH* in the region between *TBPH* and the nearby gene *bgcn*. We examined the precise sequence where both the 5’ and 3’gRNAs were designed to target in this sequence. We found that the donor vector had inserted into the genome at the location where a DSB was designed to have been induced by one of the gRNAs. At this location, on both sides of the insertion, the PAM sequence of the target region was modified to match the edited PAM sequence of the template donor plasmid consistent with this being the insertion point of the vector sequence. From this we conclude that the entire template vector has inserted into the genome at the site where a DSB was designed to be induced by one of the gRNAs. To determine if the HDR donor vector sequence was present in all incorrectly edited animals, we designed primers to amplify the region between the lac promoter and pUC ori sequence of pCR4. Using these primers, we found that 100% (4 out of 4) of incorrectly edited animals, i.e. where *Drosophila TBPH* and *hTDP-43* were both expressed, had pCR4 sequences inserted into their genome. No product could be amplified from the genomic DNA derived from background control or correctly edited animals (Figure 2F). Thus, in these anomalously edited animals, we observed an undesired incorporation of the entire HDR donor vector into the genome. We also observed that the undesired template vector insertion could be inherited stably in animals across multiple generations consistent with germline genome insertion.

### Unwanted HDR vector genomic incorporation during insertion of sequences into an endogenous gene

We wondered if the high rate of insertion of vector sequences into the genome during *TBPH* editing was associated with the particular gRNAs we had utilised for this HGR editing procedure, or due to the nature of the large editing event that we had attempted to achieve. To investigate this further, we examined a different type of editing event, i.e. in a different target gene and employing different gRNAs. The Vesicular GABA transporter (*VGAT*) (26) is required for the release of the neurotransmitter GABA. We wished to insert an ALFA epitope tag (27) in frame into the *VGAT* ORF to enable analysis of protein expression. Again, we designed two gRNAs using CRISPR Optimal Target Finder and an HDR donor plasmid consisting of homology arms, *VGAT* sequence that included the epitope tag and an eye DsRed selection cassette, but employed a different cloning vector, pHSG298 (Figure 3A) guided by a prior studies (28). Fifteen genome edited animals were recovered with eye DsRed expression and the presence of the ALFA tag was assessed by PCR genotyping. Using primers specific for the ALFA tag and *VGAT* flanking regions outside of the region of HDR homology arms, we confirmed the presence of an ALFA tag insertion in all lines. We then tested how many of these lines had undesired insertion of pHSG298 sequences. We found that 10 lines (66.6%) had incorporation of pHSG298 sequences into the genome, with only 5 edited (33.3%) lines showing no evidence of a donor vector insertional event (Figure 3B,C). Importantly, we examined the lines with undesired vector incorporation using immunohistochemistry and western blotting to confirm correct ALFA tag insertion into *VGAT*. We confirmed that indeed several lines did have ALFA tag insertion into the *VGAT* gene (data not shown). Therefore, unlike TPBH gene replacement described above, correct *VGAT* genome editing could occur in tandem with the unwanted insertion of the HDR template vector sequence into the genome. Our results also indicated that the choice of HDR donor vector does not seem to be critical, as pHSG298 sequences were inserted similarly to pCR4. Thus, in an independent HDR based genome edit of an endogenous *Drosophila* gene, we also observed digressive template vector genomic incorporation.

**Figure 3.**
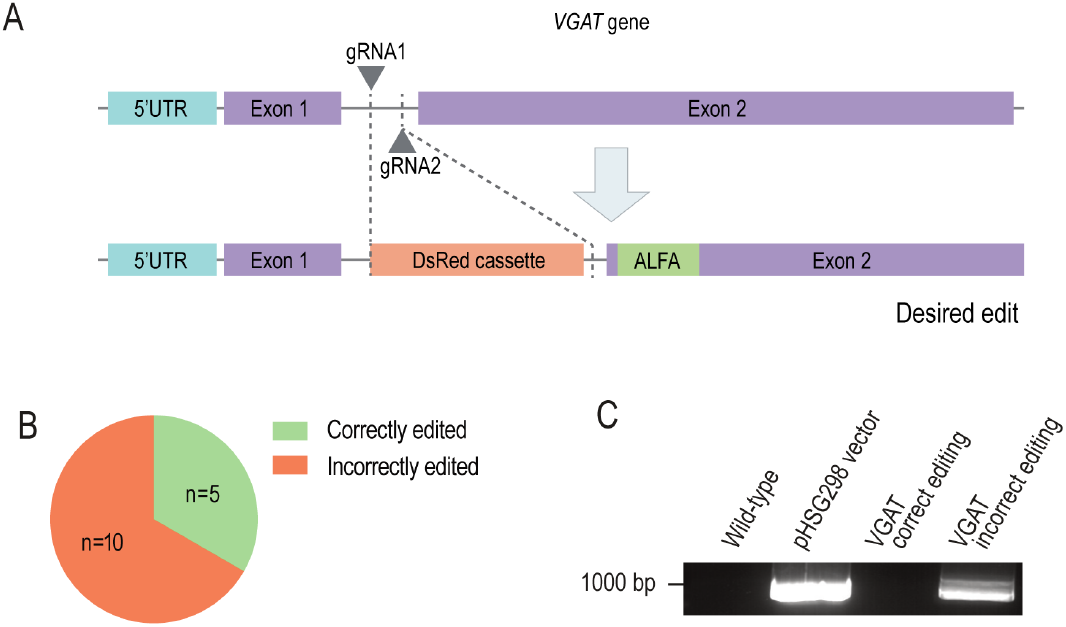
Undesired template vector integration in genome editing to insert an epitope tag. (A) Strategy for CRISPR/Cas9 plus HDR insertion of an ALFA tag in frame in exon 2 in the *VGAT* gene. Two gRNAs enable two DSBs between exons 1 and 2 of the *VGAT*. The ALFA tag and a removable DsRed cassette, flanked by ∼1 kb long homology arms, in a pHSG298 vector acts as a template for HDR. Correct editing results in a DsRed cassette in intron 1 and an in-frame ALFA tag sequence within exon 2. (B) Chart of the ratio of correctly inserted tagged *VGAT Drosophila* lines, compared to those with inserted template vector sequence (C) Image of an agarose gel of products amplified using PCR using primers specific for the vector pHSG298. pHSG298 sequences can be amplified from pHSG298 and genomic DNA derived from un-wanted editing of *VGAT Drosophila* lines, but not correctly edited *VGAT Drosophila* lines or controls. Diagrams are not to scale.

### HDR template vector mutagenic insertion editing a viable gene

Both *TBPH* and *VGAT* are recessive lethal when mutated (29, 30). We wondered, even though selection for editing events is carried out in heterozygotes, if this could in some way bias towards undesired HDR template vector genome incorporation. Also, both of the genome edits described above included insertion of exogenous sequences (human *hTDP-43*, ALFA epitope tag) into the genome. We therefore examined another HDR editing event, where our aim was to mutate nucleotides within the Glutamate Receptor IIA (*GluRIIA*) gene, which is homozygous viable even in protein null mutants (31). Similar to the above genome editing strategies, we designed two gRNAs to target exon 12 of *GluRIIA* and an HDR vector (in pCR4) with homology arms designed to replace two amino acids in *GluRIIA* with modified variants, again using eye DsRed expression to select editing events (Figure 4A). Twelve DsRed expressing *Drosophila* lines were recovered and were analysed by PCR genomic amplification and sequencing. Of these 12 lines, only one had the correct edit, as confirmed by DNA sequencing (Figure 4B). From the other 11 lines, DNA sequencing showed two peaks in the sequencing chromatogram, revealing the presence of the unmutated *GluRIIA* nucleotides in addition to the desired modified nucleotides. This suggested that both variants of this exon sequence were present in the genome of these animals. To confirm that this was due to insertion of the HDR vector into the genome, we carried out a PCR for pCR4 vector sequences from genomic DNA derived from these incorrectly edited animals (Figure 4C). All 11 incorrectly edited lines were confirmed to have pCR4 sequences inserted into their genome. Thus, again, with a third set of independent gRNAs targeting another independent gene, we observed a high frequency of anomalous template donor vector insertion into the genome as a problematic consequence of HDR based genome editing.

**Figure 4.**
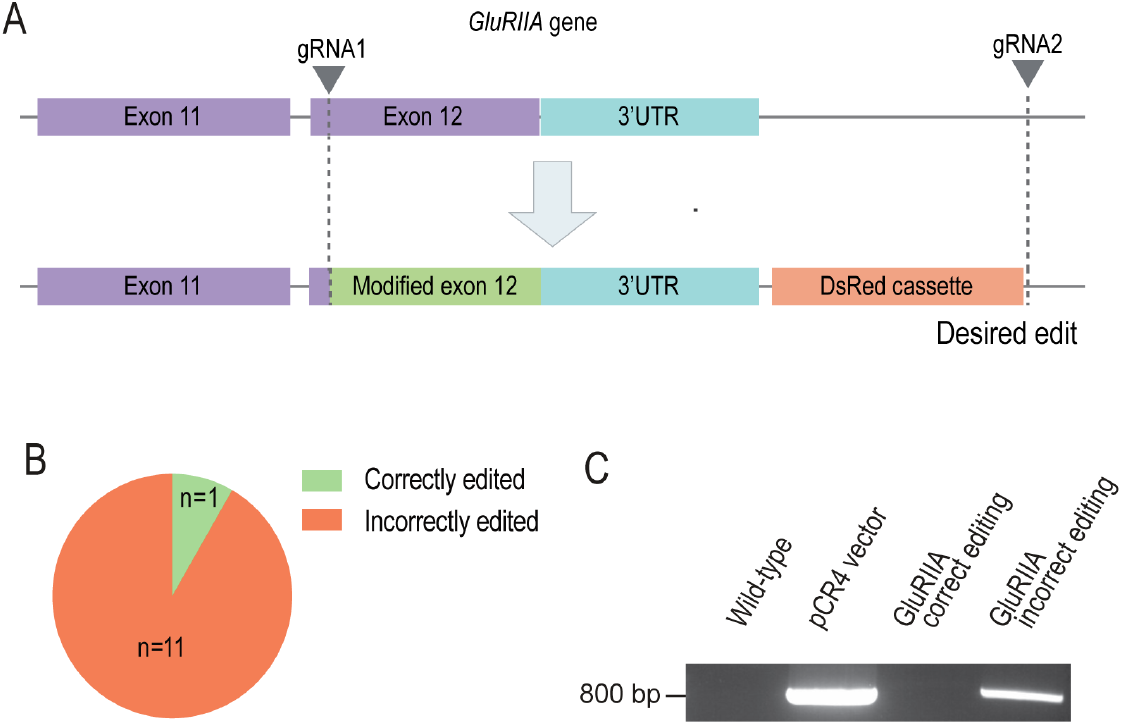
Undesired HDR vector genomic integration in genome editing to alter single nucleotides. Illustration of the CRISPR/Cas9 plus HDR editing strategy to edit single nucleotides in exon 12 of the *GluRIIA* gene by employing two gRNAs to enable DSBs. A pCR4 plasmid containing the mutated *GluRIIA* exon 12 and a removable DsRed cassette, flanked by ∼1 kb long homology arms was provided a template for HDR. Correct editing results in a modified version of exon 12 labelled by a selectable DsRed cassette. (B) Chart illustrating the number of lines with correct exon replacement (n=1) compared to the number of lines with incorrect template vector backbone insertion (n=11). (C) Agarose gel image of genomic DNA PCR amplification using primers specific for pCR4. The vector backbone sequence was identified in incorrectly edited *GluRIIA Drosophila* lines, but not in the correctly edited *GluRIIA Drosophila* line or controls. Diagrams are not to scale.

### Undesired insertion of HDR template vector can be excluded through use of a selectable marker

Selection markers are widely used in yeast, mice and *Drosophila* to detect exogenous sequence insertional events (13–15, 32, 33). As we wished to continue to use a DsRed piggyBAC selection cassette (22) to identify successful editing events (and subsequently ‘scarlessly’ remove it after successful edit identification), we sought to incorporate an additional selection marker into the HDR template backbone to identify and exclude vector insertional mutagenic events from subsequent analysis. Mini-white (34) is a commonly used selectable marker in *Drosophila*, and in addition to eyes absent selection, has previously been evaluated as a template vector negative selection marker (35). However mini-white selection proved unreliable for detection of vector genomic insertion in our hands (data not shown). We therefore decided to design a vector using a completely exogenous gene for negative selection with a bright selection marker that would be easy and quick to score. We designed a HDR template cloning vector to produce expression of ZsGreen1 (36), a very bright green fluorescent protein, under the eye P3 promoter (37) only if the vector backbone becomes incorporated into the genome. We dubbed this vector pVID - Vector Insertion Detector (Figure 5A).

**Figure 5.**
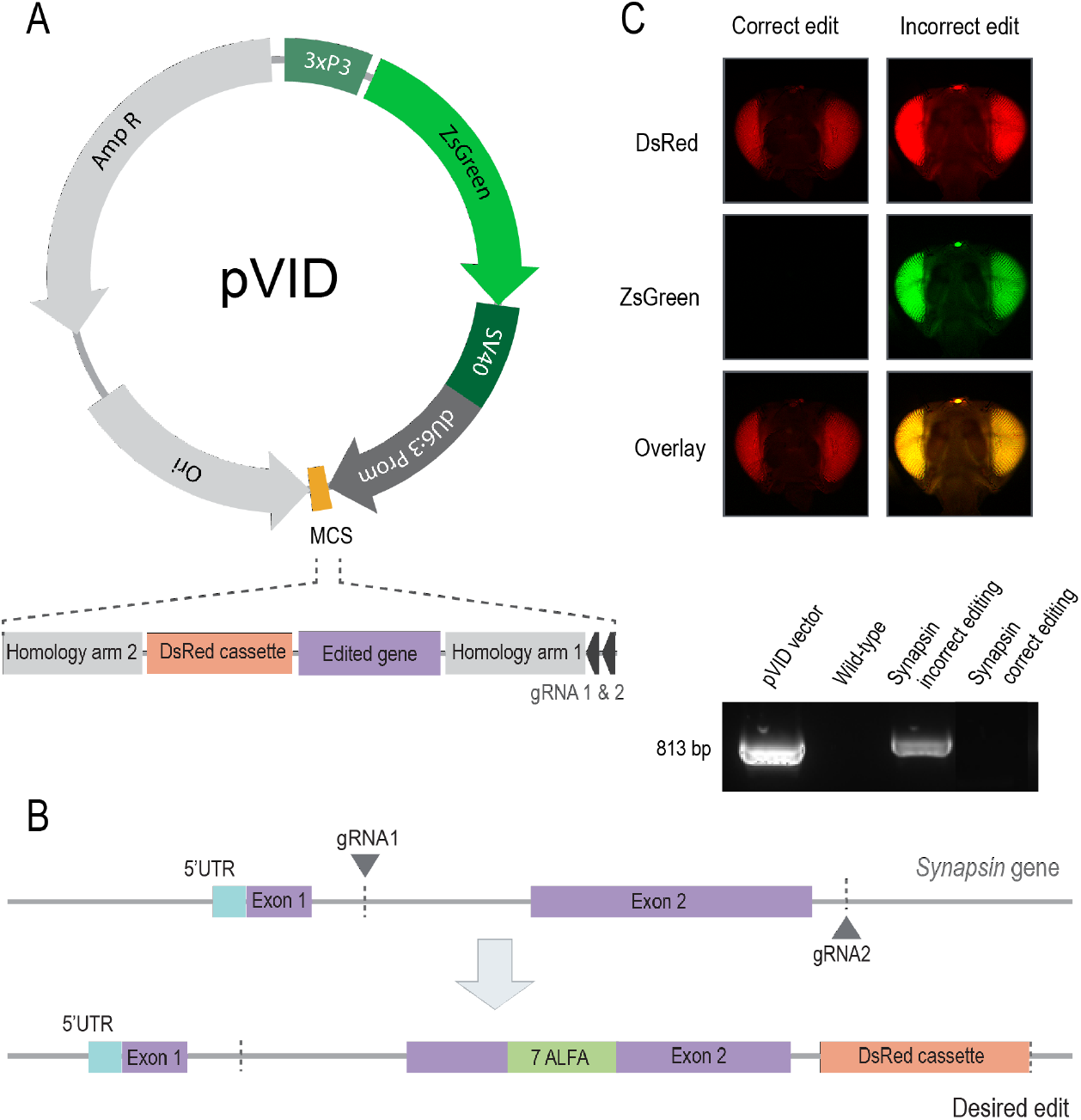
The pVID HDR template vector enables the identification of *Drosophila* genome edited lines with undesired vector integration events using ZsGreen. A) Graphical map of pVID vector (to scale). It includes an Ampicillin resistance, a *Drosophila* U6:3 promotor for gRNA expression (dark grey), a multiple cloning site (MCS) for the insertion of the HDR template sequence (orange) and a ZsGreen selection cassette (green) with its 3xP3 eye promoter, ZsGreen fluorescent protein coding sequence and SV40 terminator. In our editing strategy, the HDR template sequence contains the desired edit and a DsRed cassette selection flanked by two homology arms, plus two gRNAs under the control of dU6:3 promoter (dark grey). B) Strategy for CRISPR/Cas9 plus HDR insertion of seven ALFA tags in frame in exon 2 in the *Synapsin* gene. Two gRNAs enable two DSBs between exons 1 and 2 and after exon 2. Correct editing results in a DsRed cassette in intron 2 and an in-frame 7ALFA tag sequence within exon 2.(C) Image of eye scoring for correct and incorrect edit. DsRed positive eye expression indicates the presence of the desired genomic edit, DsRed and ZsGreen eye expression indicate that edit is accompanied by an undesired vector backbone insertion. (D) Agarose gel image of pVID vector backbone PCR amplification from edited *Synapsin Drosophila* lines and controls, showing undesired vector insertion within the genome of ZsGreen expressing animals. Diagrams are not to scale.

To evaluate the effectiveness of pVID, we performed genome editing of *Synapsin* (38), which encodes a phosphoprotein involved in vesicle release, designed to insert ALFA epitope tags within the *Synapsin* open reading frame. Similar to the genome editing procedure described above, we designed two gRNAs using CRISPR Optimal Target Finder and an HDR donor plasmid consisting of homology arms, a *Synapsin* sequence that included the epitope tags and an eye DsRed selection cassette, but employed pVID as the cloning vector (Figure 5B). 16 DsRed expressing *Drosophila* lines were recovered. Of these, 8 (50%) also had ZsGreen expression. We carried out PCR for pVID vector sequences from genomic DNA derived from ZsGreen expressing lines and found all (8 of 8) had template vector insertion. In contrast, PCR for pVID vector sequences from genomic DNA derived from DsRed expressing lines which did not have ZsGreen expression, showed no evidence of vector template sequences (8 of 8), plus all had the expected *Synapsin* genome edit. We additionally utilised pVID for second CRISPR/Cas9 HDR genome editing procedure to insert sequences into a different gene, GluRIIC, which encodes an obligate Glutamate receptor (39). The GluRIIC edit included a HDR template with the desired insertion and an eye DsRed selection cassette. As was the case for *Synapsin* editing, we recovered lines with both DsRed and ZsGreen eye expression (5 of 6) and a line with DsRed eye expression but no ZsGreen eye expression (1 of 6). We carried out PCR for pVID vector sequences from genomic DNA derived from ZsGreen expressing lines and found all (5 of 5) had HDR template vector insertion. In contrast, the line which only expressed DsRed had no evidence of HDR template vector insertion. We conclude that use of pVID as the backbone vector for HDR templates utilised for genome editing allows the successful identification and exclusion of undesired mutagenic HDR vector insertion events.

## Discussion

CRISPR/Cas9 based genome editing has been widely deployed to introduce, remove or modify genetic sequences in many model organisms including *Drosophila* (10, 23, 40). Here we describe the highly frequent undesired insertion of the donor template vector into the genome in experiments designed to alter genes based upon homology directed repair. We observed this unwanted insertional mutagenesis in five fully independent HDR procedures utilising *Drosophila* germline genome editing. Looking across the totality of the genome edits described here, an average of 54% (n=71) of the edited lines we generated had undesired vector insertion, consistent with other reports (17). The procedures we employed to design our HDR edits were not unusual. gRNA sequences were designed using CRISPR Optimal Target Finder based on conventional on-target and off target selection criteria (21) (Table 1). The germline Cas9 expressing animals we employed are also widely used (25). Our HDR repair templates utilised conventional design with ∼1 kb flanking homology arms together with the sequence to be inserted or edited including mutated PAM sequences (9). Slightly less common, yet also widely employed, was our use of an eye expression DsRed selection cassette to identify editing events, thanks to the ability to ‘scarlessly’ remove it after successful transgene or genome edit identification (22). Indeed, the presence of DsRed allowed us to unambiguously identify events where the genome was edited. HDR donor vectors that do not include a visible maker also integrate into the genome at high frequencies but are much harder to detect (17). The precise sequence of the HDR vector also does not seem to influence genome integration as we observed insertion of both pCR4 (24), pHSG298 (41) and, as described here, pVID.

**Table 1.**
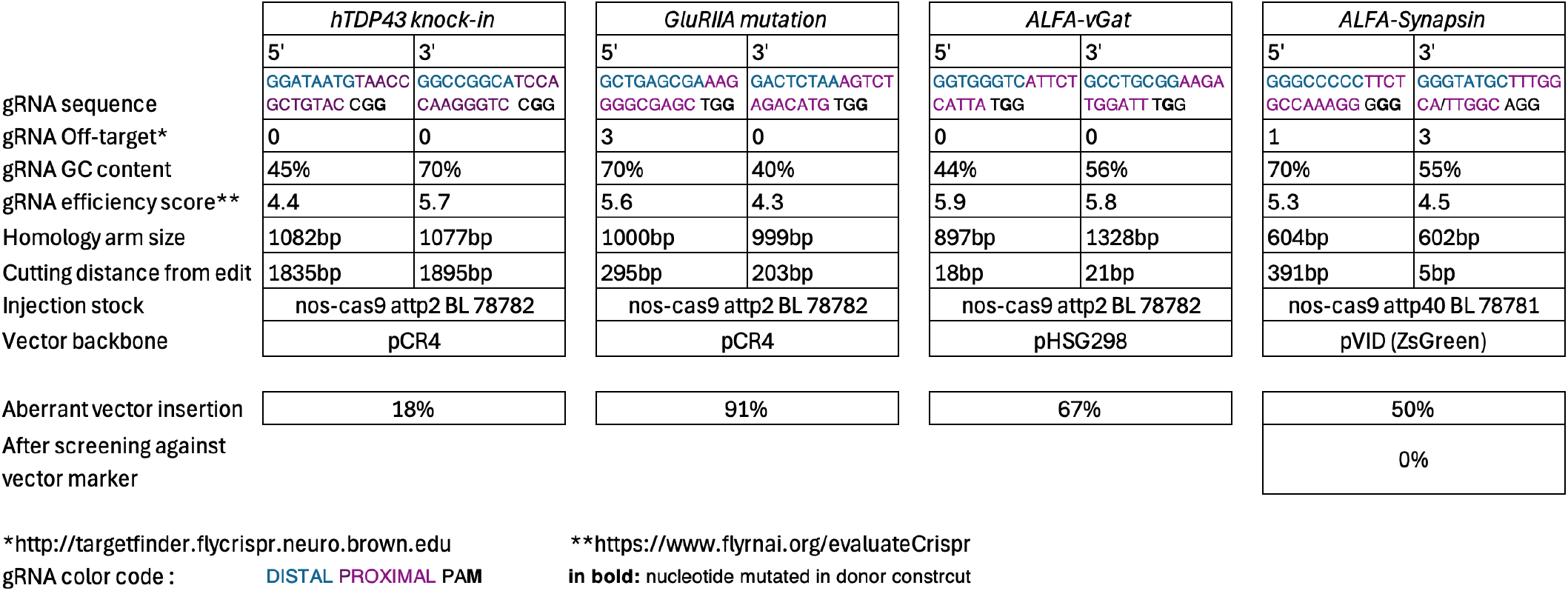
Information about the characteristics of the guide RNAs used.

Integration of the HDR template is a major issue for subsequent analysis of edited animals, potentially giving rise to misleading results. For example, if the genome edit has not occurred as designed but is falsely indicated as correct, by genomic PCR from the integrated vector, results may be misinterpreted. Alternatively, mutagenic integration of the vector sequence may cause phenotypes not attributable to the designed edit. Unexpected insertions have previously been observed during CRISPR/Cas9 based mouse genome editing, at a rate of 10% when targeting individual genomic sites with more than one gRNA, compared to 4% when targeting individual genomic sites with a single gRNA (12). In our experiments, we speculate that one of the gRNAs we designed to target a gene has a lower efficacy leading recombination to only occur at one gRNA site instead of two, causing a single homologous recombination event and insertion of the vector template into the locus. Indeed, it has been reported that gRNAs can have differing levels of efficiency which can affect editing throughput (42, 43). It is possible, but untested by us, that HDR strategies based upon the use of a single guide RNA might reduce undesired HDR donor vector insertion (17, 44) though perhaps at the cost of efficiency (45).

It is also possible that other factors, besides the efficiency of the gRNAs, enable undesired vector backbone insertion. One possibility is that the HDR vector template becomes linearised at some frequency which could facilitate insertion into the genome. Linearized templates are becoming increasingly employed in the field of CRISPR/Cas9 in several models, due to increased efficiency of editing (44, 46, 47). However, it is noteworthy that during HDR germline editing of mice, insertion of linear HDR template sequences has also been observed, not just in one copy but in several head-to-tail copies (48). These insertions were shown to be produced by both HDR and NHEJ, or in some cases, a combination of both mechanisms. Similar mechanisms could be compatible with the insertion events that we observe. Therefore, both with linear and circular templates, it is important to remain vigilant by including controls for undesired vector template insertions. Consistent in all of our experiments is also the fact that we express Cas9 in the *Drosophila* germ cells under the nos promoter to enable HDR to occur. It is possible that other Cas9 sources, such as Cas9 driven by a different promoter, or at a lower or higher expression level, could influence the insertion of the backbone vector though very high levels of Cas9 can be toxic (49).

In order to improve efficiency of HDR-mediated genome editing in our laboratory, we now routinely use two selection cassettes, a positive selection (subsequently removable) dsRed cassette to indicate an editing event has occurred and a negative selection ZsGreen cassette to eliminate events where the HDR template vector has inserted into the genome. We would suggest that others using similar approaches might consider the same, which unfortunately, since HDR template vector insertion is as we describe common, requires the generation of many additional lines to select only those that have the desired edit. Ultimately however, it would be desirable to understand more about the factors that lead to undesired HDR vector genomic integration, in order to design strategies to avoid this unwanted, labor adding and detrimental outcome of CRISPR/Cas9 based HDR dependent genome editing.

## Limitations of the Study

We note that while we observed the insertion of the template vector using three different vector template backbones, there were still similarities between the five different *Drosophila* lines we analysed in this paper. For example, in all lines, two gRNAs and a removable DsRed selection cassette were employed. In future experiments it would be beneficial to investigate the possible presence of template vector in lines which have been produced by a different number of gRNAs using a different selection marker. We also tested for undesired insertional mutagenesis only during *Drosophila* melanogaster germline genome editing and not in other model organisms or cell lines where, given similar procedures, this undesired outcome could also potentially occur. In addition, we used exclusively the nos promoter to highly express Cas9 in *Drosophila* germ cells, and it is possible that the Cas9 concentration could affect vector template integration.

## Acknowledgements

We would like to thank Dr. Steven Stowers for assistance with vectors and the design of the *VGAT* targeting strategy plus for critical reading of the manuscript. We also are grateful to other members of the McCabe laboratory for their criticisms. EK was funded by the European Union’s Horizon 2020 research and innovation program under the Marie Skłodowska-Curie grant agreement No 721802. MR was funded by the European Union’s Horizon 2020 research and innovation program under the Marie Skłodowska-Curie grant agreement No 945363. This work was supported by the Swiss National Science Foundation grant numbers: CRSII3_154455, 31003A_179587 and 320030-232324 to B.M.

## Author Contributions

Conceptualization, B.M.; Investigation, E.K., E.R, M.R, J.A.; Writing – Original Draft, E.K. and B.M.; Writing – Review & Editing, E.K., B.M., E.R., M.R.; Funding acquisition, B.S. and B.M.

## Materials and Methods

### Plasmid construction

For all plasmids, homology arms, the desired edits, and the selection cassettes were PCR-amplified using a high-fidelity DNA polymerase (Platinum SuperFi II, Invitrogen) and assembled via HiFi DNA Assembly (Invitrogen), unless stated otherwise. All gRNAs were chosen using CRISPR Optimal Target Finder (21).

### hTDP-43 knock-in HDR construct in pCR4

The full assembly was inserted by Topo cloning into pCR4. The two gRNAs cloned each in a pCDF3 vector (gRNA sequences in Table 1) targeted neighbouring genes cpes and bgcn, while making sure to not change the amino acid sequence of either gene. The inserted sequence included both *TBPH* 5’ and 3’UTRs but with the *TBPH* coding sequence replaced by a *Drosophila* codon optimised version of the human *hTDP-43* cDNA. The whole edit was surrounded by ∼1 kb homology arms.

### ALFA-VGAT tagged HDR construct in pHSG298

The cloning strategy was based on previous vGat genome editing (28). The full assembly was inserted by restriction digestion - ligation into pHSG298. The two gRNAs cloned in pCFD4 vector (gRNA sequences in Table 1) targeted the intronic region between *VGAT* exon 1 and exon 2. The insertion included an ALFA tag as well as a DsRed selection cassette. The whole edit was surrounded by ∼1 kb homology arms.

### GluRIIA exon HDR construct in pCR4

The full assembly was inserted by Topo cloning into pCR4. The two gRNAs cloned each in a pCDF3 vector targeted the beginning of exon 12 and the intergenic region between *GluRIIA* and *GluRIIB*. (gRNA sequences in Table 1) The inserted sequence included another version of exon 12 followed by the endogenous 3’UTR and a DsRed selection cassette. The whole edit was surrounded by ∼1 kb homology arms.

### 7ALFA-Synapsin HDR construct in pVID

The 5’ gRNA targeted the intronic region between exon 1 and 2 of *Synapsin*, and the 3’ gRNA targeted the intronic region between exon 2 and 3 (gRNA sequences in Table 1). The edit included 7 copies of ALFA tag and an excisable dsRed selection cassette. Upstream of the edit was two gRNAs with their respective scaffold sequence, interspaced by a tRNA sequence (50, 51). The edit was flanked by two 600-700bp Homology arms. The full sequences were synthesized and cloned in a pVID vector by Genscript.

### GluRIIC HDR Construct in pVID

The 5’ gRNA targeted the middle of exon 10, and the 3’ gRNA targeted the intronic region between exon 2 and 3 (gRNA sequences in Table 1). The edit included a Myc tag followed by an excisable dsRed selection cassette. Upstream of the edit was two gRNAs with their respective scaffold sequence, interspaced by a tRNA sequence (50, 51). The edit was flanked by two 600-700bp Homology arms. The full sequences were synthesized and cloned in a pVID vector by Genscript.

### pVID HDR vector generation

A *Drosophila* eye negative selection cassette composed of a 3xP3 promoter (48), a ZsGreen fluorescent protein (52, 53) and a SV40 terminator was inserted upstream of a dU6:3 gRNA promotor. Downstream of the promoter a multiple cloning site (MCS) was introduced to allow simple insertion, by restriction digestion and ligation, of the specific gRNAs and the HDR editing template to produce an ‘all-in-one’ CRISPR genome editing construct. The region including the green negative selection cassette and MCS was synthetized and subcloned into pUC57-mini vector by Genscript. The pVID full sequence is available with Genbank (54) accession: PV446458. Addgene 236635.

### Generation of *Drosophila melanogaste*r lines

To generate the transgenic lines, *Drosophila melanogaster* embryos expressing Cas9, under the germ cell specific nos promoter, were injected by either Genetivision (Stafford, TX, USA) or FlyORF (Zürich, Switzerland) with the HDR construct and if required the appropriate gRNA expressing vectors.

### Immunoblotting

20 adult *Drosophila* heads were lysed in RIPA lysis buffer (140 mM NaCl, 1 mM EDTA, 1%Triton X-100, 0.5 mM EGTA, 10 mM Tris-Cl pH 8, 0.1% SDS) by homogenisation with a pestle and sonication for 5 s. The samples were subsequently centrifuged for 1 min at 12 000 x g, the supernatant collected, and the protein concentration measured with a Micro BCA Protein Assay kit (Thermo Scientific). Before loading 30 µg of protein into a 4-12% Tris-Glycine gel (Invitrogen), the samples were mixed with 4x Laemmli buffer and heated for 2 minutes at 85° C. After running the gel, the protein bands were dry transferred to a PVDF membrane (iblot transfer, Invitrogen) which was blocked for 1 h in 5% milk blocking solution. The membrane was then incubated at 4°C overnight in a mix of blocking solution and primary antibodies. The primary antibodies used were anti-hTDP-43 (H00023435-M01, Novus Biologicals, 1:500), anti-TBPH (custom made, Pierce, 1:500), anti-α-tubulin (T6199, Sigma, 1:10 000). Membranes were incubated with secondary antibodies (goat anti-rabbit, 111-035-003, Jackson ImmunoResearch, 1:10 000) (goat anti-mouse 115-035-003, Jackson ImmunoResearch, 1:10 000) diluted in blocking solution for 1 h. Chemiluminescent substrate (WesternBright Sirius, Advansta) was added to enable imaging. To stain with several consecutive antibodies the membranes were stripped for 10 min at 37°C (Restore Western Blot Stripping Buffer, Thermo Scientific) and re-blocked.

### PCR of region between incorrect insert and endogenous gene

The genomic DNA of all *Drosophila* lines were purified using a commercial kit (DNeasy Blood and Tissue kit, Qiagen). To detect the DNA sequence between the *hTDP-43* gene and the *TBPH* gene in the incorrectly inserted *hTDP-43* knock-in *Drosophila*, we performed a PCR using a proofreading DNA polymerase (Platinum SuperFi II, Invitrogen), a reverse primer in the 5’ end of the *hTDP-43* gene (TTCTCATCCTCG-GTCACGCG) and a forward primer in the 3’ end of the *TBPH* gene (GGAACACCGGTTGGTCGAAC). The PCR product was run on an agarose gel and the band was purified using a gel extraction kit (GeneJET Gel Extraction Kit, Thermo Scientific) and a reaction clean-up kit (MinElute Reaction cleanup kit, Qiagen). The PCR product was then cloned into a pCR8 vector with pCR™8/GW/TOPO™ TA Cloning Kit (Invitrogen) and sequenced.

### Identification of the pCR4 or pHSG298 template vector backbones in the *Drosophila* genome

To identify the presence of the vector backbone in the *Drosophila* genome, we purified the genomic DNA using DNeasy Blood and Tissue kit (Qiagen), and performed a PCR using primers within the lac promoter and pUC ori in the pCR4 or pHSG298 plasmid back-bone (forward: CGCGTAATCTGCTGCTTGCAA, reverse: TGTCGTGCCAGCTGCATTAATG). The resulting band should be 817 bp long in the pCR4 backbone plasmid, and 1001 bp long in the pHSG298 plasmid backbone. For pVID vector backbone, we performed a PCR using primer covering the region between the zsGreen CDS and the dU6:3 promoter (forward: 5’-GACAGACAACTGGGAGCCTAGTTG-3’, reverse: 5’-TGAGTAGGTGGCGTTTCATTCTACTC-3’). The resulting band should be 813bp long.

### Identifying undesired HDR template vector genomic insertion with pVID

To select only animals where editing has occurred but the template HDR vector has not become incorporated into the genome we selected for DsRed expressing eye fluorescence and subsequently ZsGreen green expressing eye fluorescence. Flies exhibiting ZsGreen eye fluorescence indicted the undesired genomic integration of the pVID vector backbone. Fluorescence screening was performed using a fluorescence stereomicroscope (MC165F Leica) equipped with a GFP and dsRed filter. Images were taken using a fluorescence CCD camera (DFC3000G, Leica).

## Key Resources Table

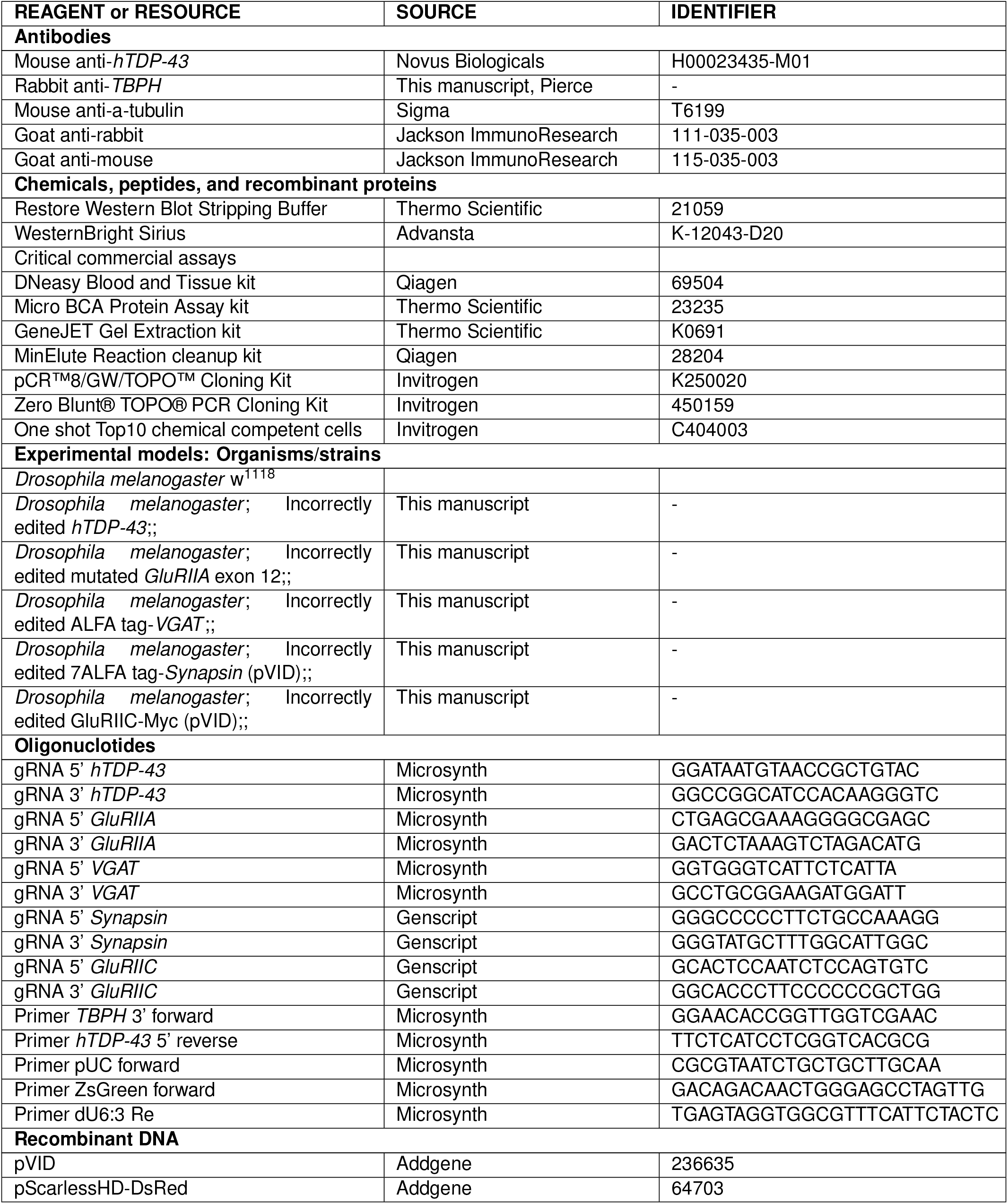

